# Rhythmic auditory stimulation rescues cognitive flexibility in mutant mice with impaired gamma synchrony

**DOI:** 10.1101/2021.11.15.468681

**Authors:** Jingcheng Shi, Aarron J. Phensy, Vikaas S. Sohal

## Abstract

Neural synchronization at gamma (∼40 Hz) frequencies is believed to contribute to brain function and be deficient in disorders including Alzheimer’s disease and schizophrenia. Gamma-frequency sensory stimulation has been proposed as a non-invasive treatment for deficient gamma synchrony and associated cognitive deficits, and has been shown to be effective in mouse models of Alzheimer’s disease. However, both the mechanism and applicability of this approach remain unclear. Here we tested this approach using mutant (*Dlx5/6*^+/-^) mice which have deficits in gamma synchrony and the ability to learn to shift between rules which use different types of cues to indicate reward locations. 40 Hz auditory stimulation rescues rule shifting in *Dlx5/6*^+/-^ mice. However, this improvement does not outlast the period of stimulation, and is not associated with normalized gamma synchrony, measured using genetically encoded voltage indicators. These results show how gamma-frequency sensory stimulation may improve behavior without fully restoring normal circuit function.

## INTRODUCTION

Gamma-frequency (30-100 Hz) synchronization can be observed in neural activity at many scales, ranging from EEG recorded outside the skull, to the phase-locked firing of individual neurons. While the significance of such activity has long been controversial (Cardin, 2016; Sohal, 2016), many recent studies have shown that synchronized gamma oscillations play key roles in cognition. For example, we have shown that gamma synchrony between prefrontal parvalbumin (PV) interneurons normally increases when mice learn to shift between rules which use different types of cues to indicate the location of a reward, and that optogenetically disrupting this synchrony is sufficient to disrupt the learning of rule shifts (Cho et al., 2020). Conversely, mutant (*Dlx5/6*^+/-^) mice which have deficits in PV interneuron gamma synchrony have deficits in rule shifting that are rescued when gamma synchrony is restored, either optogenetically or pharmacologically (Cho et al., 2020, 2015). Other studies have shown that optogenetically enhancing gamma oscillations can improve attention (Kim et al., 2016), modulate sensory detection (Siegle et al., 2014), and enhance memory consolidation (Kanta et al., 2019). These studies and others (Marissal et al., 2018) support the idea that normal gamma synchronization contributes to cognition, such that restoring deficient gamma synchrony may ameliorate cognitive deficits associated with conditions such as schizophrenia (Gonzalez-Burgos et al., 2015; Uhlhaas and Singer, 2010).

Recently, there has been great interest in the idea that deficits in gamma synchrony may contribute to the pathophysiology of Alzheimer’s disease, such that enhancing gamma synchrony might be therapeutic (Adaikkan et al., 2019; Iaccarino et al., 2016; Liu et al., 2021; Martinez-Losa et al., 2018; Martorell et al., 2019; Verret et al., 2012). A particularly exciting possibility is that non-invasive rhythmic sensory stimulation (gamma entrainment using sensory stimulation or ‘GENUS’) might entrain neural activity across many regions, leading to decreases in Alzheimer’s pathology and improvements in cognition, at least in mouse models (Adaikkan et al., 2019; Iaccarino et al., 2016; Martorell et al., 2019). While such stimulation has been shown to lead to decreases in measures of cellular pathology and improvements in behavior, the exact mechanism through which it operates and extent to which it might be generally applicable remain unclear.

Here we sought to explore these issues using a task, mouse model, and measures of gamma synchrony we have previously characterized (Cho et al., 2020, 2015). We previously described a method to measure synchrony between parvalbumin (PV) interneurons in the left and right medial prefrontal cortex (mPFC) using genetically encoded voltage indicators. Using this method, we showed that when mice shift from an initially learned rule to a new rule which uses cues that were previously present but irrelevant to trial outcomes, cross-hemispheric gamma-frequency (∼40 Hz) synchrony between PV interneurons normally increases after error trials (when mice follow the original rule but fail to receive an expected reward). In *Dlx5/6*^+/-^ mice, this increase in gamma synchrony following error trials is lost and rule shift learning is severely impaired. However, restoring gamma synchrony by delivering in-phase 40 Hz optogenetic stimulation to PV interneurons in-phase across both hemispheres normalizes rule shifting in *Dlx5/6*^+/-^ mice. Low (sub-sedative and sub-anxiolytic) doses of the benzodiazepine clonazepam (CLZ) also normalize performance and restore increases in PV interneuron gamma synchrony after errors during rule shifts. Notably, these effects are persistent, i.e., following one session of 40 Hz optogenetic stimulation, *Dlx5/6*^+/-^ mice will continue to perform normally weeks later, in the absence of additional stimulation. Thus, here we asked three questions: (1) Would gamma-frequency (40 Hz) sensory stimulation normalize rule shifting performance in *Dlx5/6*^+/-^ mice? (2) Would gamma-frequency (40 Hz) sensory stimulation restore increases in gamma synchrony following rule shift errors in *Dlx5/6*^+/-^ mice? (3) Would therapeutic effects of gamma-frequency (40 Hz) sensory stimulation persist after the cessation of stimulation?

## RESULTS

### GENUS rescues rule shifting deficits in *Dlx5/6*^+/-^ mice

To test whether GENUS is effective in rescuing behavioral deficits in *Dlx5/6*^+/-^ mice, we used a previously-described rule shifting task (Cho et al., 2015). The task comprises two phases: learning of an *initial association* followed by the *rule shift*. On each trial, mice are presented with two bowls containing different odor and texture cues, and they choose to dig in one bowl to find a food reward (Fig. 1a). During the initial association phase of the task, mice learn an association between one of these cues (e.g., texture A) and reward. After successful learning (indicated by reaching a criterion of 8 correct out of 10 consecutive trials), mice learn a rule shift in which a cue from the other (previously-irrelevant) modality (e.g., odor 2) now becomes associated with reward.

**Fig. 1.**
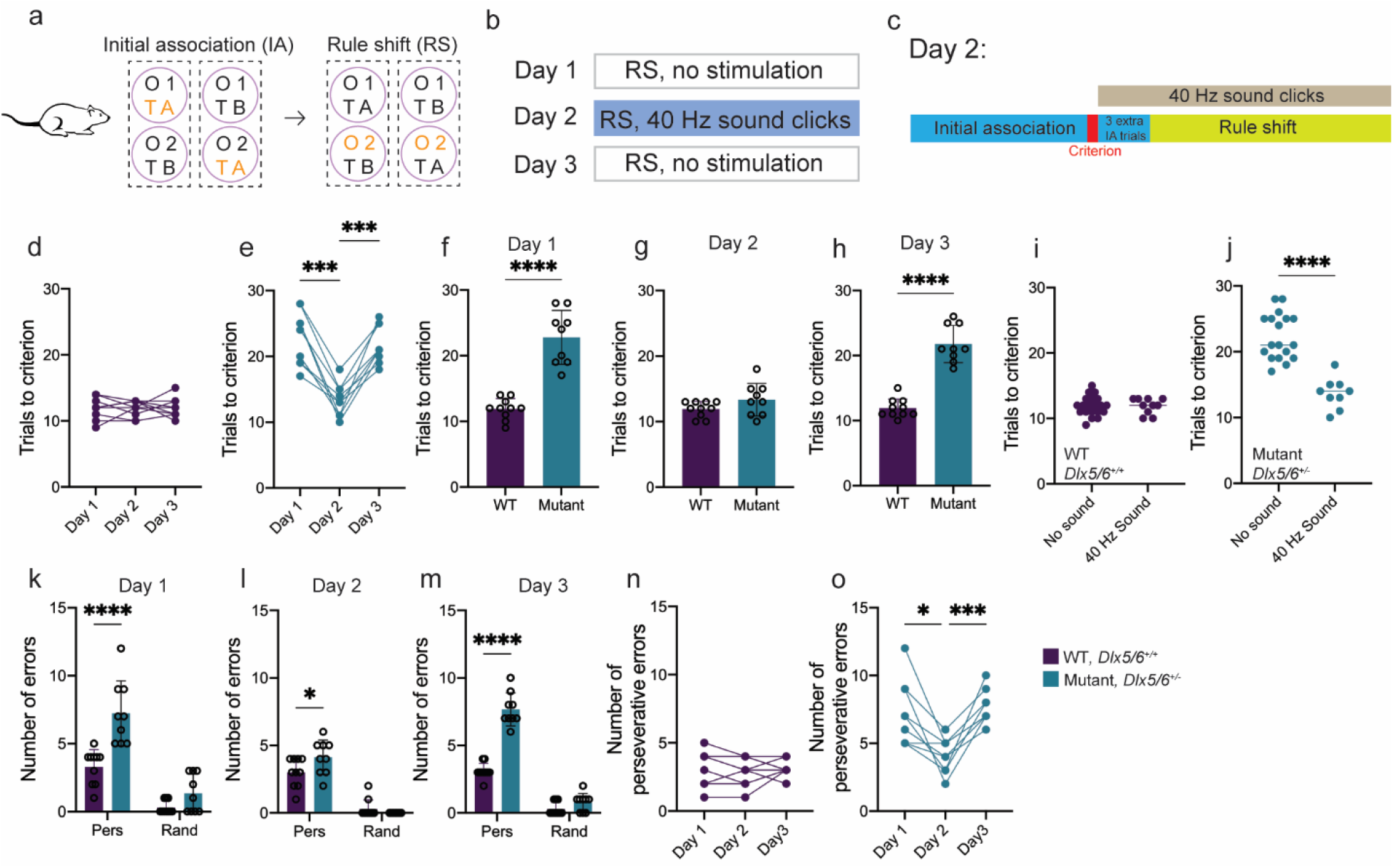
GENUS rescues rule shifting performance in *Dlx5/6*^+/-^ mice. **a**, Schematic showing the rule shift task. On each trial, mice are presented with two bowls, each with a different odor (O1 and O2) and a different textured medium (TA or TB). Mice first learn an initial association (IA) between the food reward and one cue (for example, TA). Then the rule undergoes an extra-dimensional rule shift (RS) to a cue (O2) from the other modality (texture to odor, in this example, or from odor to texture). **b**, Schematic showing the three-day rule shift paradigm. Day 1, no stimulation; Day 2, 40 Hz auditory stimulation (GENUS); Day 3, no stimulation. **c**, Schematic illustrating the delivery of stimulation on Day 2. Once mice learn the IA criterion, 40 Hz sound clicks start playing from a speaker directly above the cage. Three additional IA trials are added, then stimulation is played continuously throughout the rule shift. **d**, GENUS has no effect on the number of trials needed to learn the RS in wildtype mice. **e**, GENUS reduces the number of trials mutant mice need to learn the RS on Day 2 (n = 9 mice; one-way repeated measure ANOVA followed by Tukey’s multiple comparisons test, main effect of treatment F_1.831,14.65_=40.0, *P=0.023; Day 1 versus Day 2: ***P=0.0003; Day 1 versus Day 3: P=0.63; Day 2 versus Day 3: ***P=0.0001). **f**, On Day 1, mutant mice require more trials to learn the RS (n=10 wild-type and 9 *Dlx5/6*^+/-^ mice; two-tailed unpaired *t*-test; t_(17)_=7.8, ****P<0.0001). **g**, GENUS rescued RS performance in mutant mice to near-wildtype levels (two-tailed unpaired *t*-test; t_(17)_=1.62, P=0.12). **h**, On Day 3, mutant mice require significant more trials to complete the RS (two-tailed unpaired *t*-test; t_(17)_=9.6, ****P<0.0001). **i**, Auditory stimulation has no effect on the performance of wild-type mice. **j**, Auditory stimulation significantly reduces the number of trials mutant mice need to learn the RS (comparison of pooled data from Days 1 and 3 to Day 2; two-tailed unpaired Welch’s *t*-test; t_(22.48)_=7.6, ****P<0.0001). **k, l, m**, numbers of perseverative (P) or random (R) errors on each day of rule shift. **k**, Mutant mice make more perseverative errors on Day 1 (two-way ANOVA followed by Šidák multiple comparison, main effect of genotype: F_1,34_=81.8, ****P<0.0001; main effect of error type: F_1,34_=25.4, ****P<0.0001; error type X genotype interaction: F_1,34_=8.6, **P=0.0059; P errors: t_(34)_=5.6, ****P<0.0001; R errors: t_(34)_=1.49, P=0.27). **l**, Day 2: mutant mice still make more perseverative errors than wildtype mice (two-way ANOVA followed by Šidák multiple comparison, main effect of genotype: F_1,34_=138.4, ****P<0.0001; main effect of error type: F_1,34_=1.96, P=0.17; error type X genotype interaction: F_1,34_=5.94, *P=0.020; P errors: t_(34)_=2.71, *P=0.021; R errors: t_(34)_=0.733, P=0.72). **m**, comparison of perseverative errors in wildtype vs. mutant mice on Day 3 (two-way ANOVA followed by Šidák multiple comparison, main effect of genotype: F_1,34_=367.8, ****P<0.0001; main effect of error type: F_1,34_=99.7, ****P<0.0001; error type X genotype interaction: F_1,34_=65.5, ****P<0.0001; P errors: t_(34)_=12.8, ****P<0.0001; R errors: t_(34)_=1.34, P=0.34). **n**, GENUS has no effect on perseverative errors in wildtype mice (one-way repeated measure ANOVA followed by Tukey’s multiple comparisons test, main effect of treatment F_1.549,13.94_=0.47, P=0.58; Day 1 versus Day 2: P=0.38; Day 1 versus Day 3: P=0.85; Day 2 versus Day 3: P=0.96). **o**, GENUS significantly reduces the number of perseverative errors in mutant mice (one-way repeated measure ANOVA followed by Tukey’s multiple comparisons test, main effect of treatment F_1.744,13.95_=14.1, ***P=0.0006; Day 1 versus Day 2: *P=0.013; Day 1 versus Day 3: P=0.83; Day 2 versus Day 3: ***P=0.0007).

We tested *Dlx5/6*^+/-^ mice and their wild-type littermates on the rule shift task over three consecutive days (Fig. 1b). The first and the third day of the rule shift were performed without GENUS auditory stimulation. On the second day, mice were exposed to continuous 40 Hz auditory stimulation (clicks) throughout the rule shift portion of the task (Fig. 1c). In our previous studies, we found that in the absence of experimental manipulations, rule shifting performance is stable over three days of testing for both wild-type and *Dlx5/6*^+/-^ mice (Cho et al., 2020, 2015).

We measured the number of trials required to reach the learning criterion during both the initial association and rule shift portions of the task. As in our previous studies (Cho et al., 2020, 2015), on Day 1, in the absence of stimulation *Dlx5/6*^+/-^ mice are significantly impaired during the learning of rule shift as compared to their wild-type littermates (Fig. 1d-f), and make an excessive number of preservative errors (errors that would have been correct based on the initial association rule) (Fig. 1k). However, when mice receive auditory stimulation on Day 2, *Dlx5/6*^+/-^ and wild-type mice learn the rule shift in a similar number of trials (Fig. 1g), and make similar numbers of perseverative errors (Fig. 1l). On Day 3, performance returned to baseline levels and was again significantly worse for *Dlx5/6*^+/-^ mice compared to wild-type mice (Fig. 1h). There was no significant difference in performance for *Dlx5/6*^+/-^ mice on Day 1 (baseline) vs. Day 3 (post-stimulation), i.e., the pro-cognitive effects of auditory stimulation evident on Day 2 did not persist.

### GENUS does not increase prefrontal gamma synchrony

To evaluate whether auditory stimulation affects cross-hemispheric PV interneuron gamma synchrony, in a subset of mice that underwent behavioral testing, we also measured signals from genetically encoded voltage indicators using ‘trans-membrane electrical measurements performed optically’ (TEMPO) (Marshall et al., 2016), together with our previously-described analysis (Cho et al., 2020). We injected AAV1-DIO-Ace2N-4AA-mNeon into the bilateral mPFC to drive Cre-dependent expression of the voltage indicator Ace2N-4AA-mNeon (‘Ace-mNeon’) in *Dlx5/6*^+/-^*;PV-Cre* or *Dlx5/6*^+/+^*;PV-Cre* mice. Mice were also injected with AAV2-Synapsin-tdTomato to provide a voltage-independent reference signal for correcting artifacts related to hemodynamic sources, fiber bending, etc. Optical fibers were implanted over the bilateral mPFC to deliver light to excite Ace-mNeon and TdTomato and measure emitted fluorescence (Fig. 2a-b). As in our previous study, we used a system of lock-in-amplifiers, LEDs, dichroic mirrors, filters, and photoreceivers to separate these light signals using both wavelength and temporal modulation frequency (Fig. 2c) (Cho et al., 2020).

**Fig. 2.**
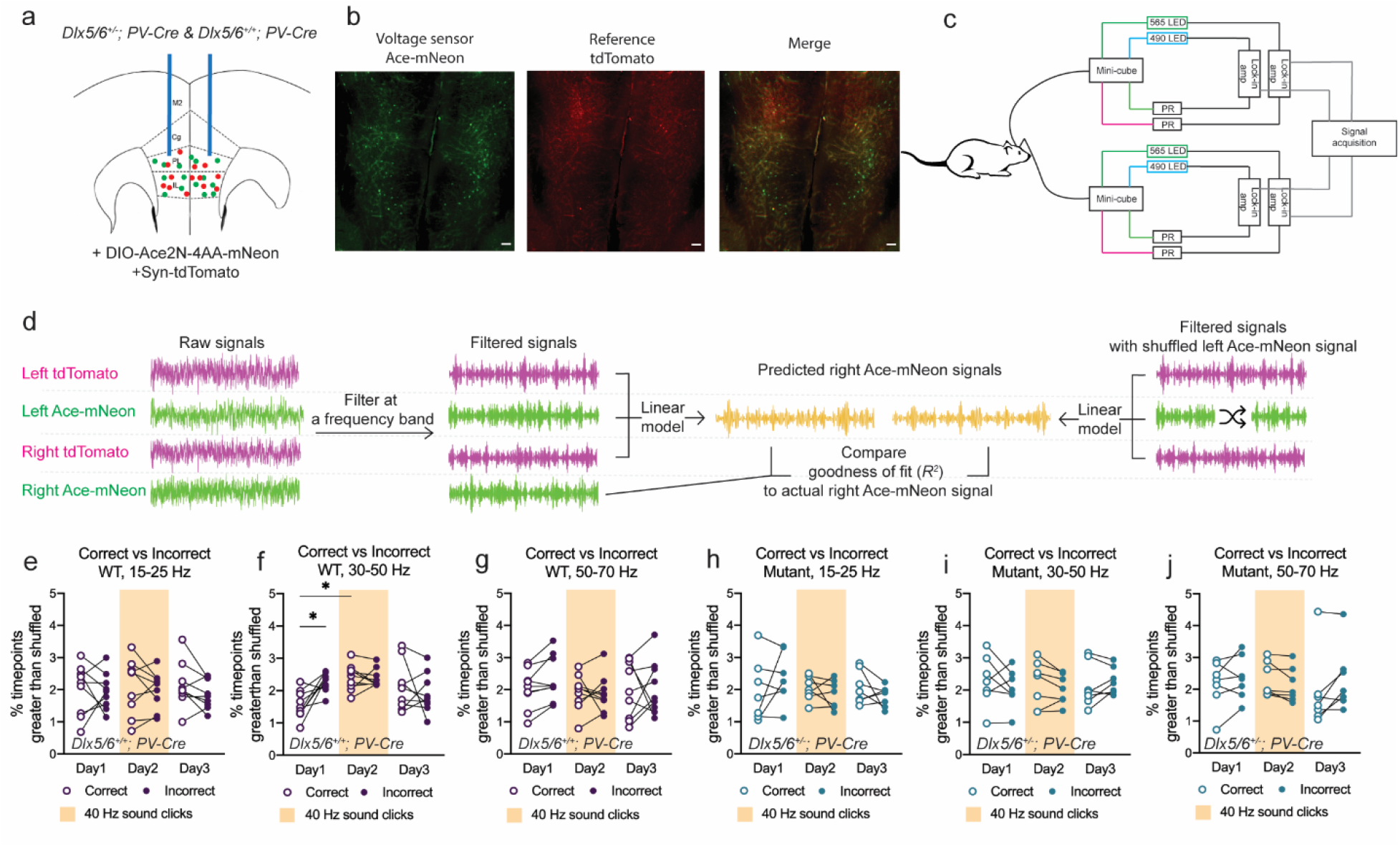
GENUS does not restore increases in cross-hemispheric PV interneuron gamma synchrony after rule shift errors in mutant mice. **a**, *Dlx5/6*^+/-^, *PV-Cre* (mutant) and *Dlx5/6*^+/+^, *PV-Cre* (wild-type) mice have bilateral AAV-DIO-Ace2N-4AA-mNeon + AAV-Syn-tdTomato injections and fiber optic implants in the mPFC. **b**, Representative image of Ace-mNeon (green, left), tdTomato (red, middle), and merged channels (right) fluorescence in coronal section of mPFC. Scale bars, 100 μm. **c**, Schematic of dual-site TEMPO recordings. Each fiber optic implant is connected by a fiber optic patch cord to a fluorescence mini-cube and delivering illumination and fluorescence to the targeted brain region. The mini-cube combines two excitation light from LEDs and separates two fluorescence to two photoreceivers (PRs). LED and PR for each fluorescence are modulated and demodulated by a lock-in amplifier. Demodulated signals from lock-in amplifiers are acquired by a multi-channel processor in real-time. **d**, Overview of TEMPO analysis. Left and right tdTomato and Ace-mNeon are filtered around a frequency of interest. Left and right tdTomato signals and the left Ace-mNeon signal are fitted to a liner model to predict the right Ace-mNeon signal. The goodness of fit (*R*^*2*^) to the real right Ace-mNeon signal is compared to that obtained using shuffled versions of the left Ace-mNeon signal to obtain the fraction of timepoints for which the prediction using the real signal significantly outperforms those using shuffled signals. **e-g**, Comparisons of the synchrony, i.e., the fraction of timepoints for which the real fit significantly exceeds that obtained using shuffled Ace-mNeon signals, following correct vs. incorrect trials during the first five rule shift trials for wildtype mice in various frequency bands. 30-50 Hz synchrony was higher following RS error trials (compared to RS correct trials) on Day 1, and higher for correct trials on Day 2 compared to correct trials on Day 1 (n=9 wild-type mice; one-way repeated measure ANOVA followed by Tukey’s multiple comparisons test, main effect of treatment F_1.787,22.30_=3.95, *P=0.023; Day 1 correct versus day 1 incorrect: *P=0.036; Day 1 correct versus day 2 correct: *P=0.016). Synchrony was not different for 15-25 or 50-70 Hz. **h-j**, Comparisons of synchrony following correct vs. incorrect trials during the first five rule shift trials for mutant mice in various frequency bands.

To measure synchrony between voltage signals from PV interneurons in the left and right mPFC, we filtered all signals in the frequency band of interest, then fit the Ace-mNeon signal on one side using both TdTomato signals (to correct for artifacts) and either the true contralateral Ace-mNeon signal, or time-shuffled versions of this signal. The fit, i.e., correlation, between two filtered Ace-mNeon signals reflects their synchrony. Thus (as in our previous study), we used the fraction of time windows in which the fit using the true Ace-mNeon signal outperformed 99% of fits achieved using shuffled versions of the Ace-mNeon signal to measure the degree of zero-phase-lag cross-hemispheric synchronization (Fig. 2d). We focused our analysis on the first five rule shift trials, which are crucial for mice to learn the new association. We compared synchrony on correct vs. incorrect trials for different frequency bands: 15-25 Hz (Fig. 2e, h), 30-50 Hz (Fig. 2f, i), and 50-70 Hz (Fig 2g, j).

Consistent with our previous studies, on Day 1, we observed that wildtype mice had significantly higher ∼40 Hz synchronization after error trials than after correct trials (Fig. 2f), whereas this was not the case for *Dlx5/6*^+/-^ mice (Fig. 2i). Surprisingly, even though we observed normal behavior in mutant mice on Day 2, GENUS did not restore the normal pattern of increased ∼40 Hz synchrony after error trials in *Dlx5/6*^+/-^ mice. Intriguingly, the pattern of ∼40 Hz synchrony was altered in wild-type mice in the presence of GENUS on Day 2. Specifically, ∼40 Hz synchrony on correct trials was significantly higher on Day 2 compared to Day 1, such that there was no longer a difference between ∼40 Hz synchrony following correct vs. error trials. This suggests that GENUS may non-specifically increase gamma synchrony in wildtype mice. Though GENUS had no lasting effect on behavior on Day 3 for either genotype, this pattern of altered gamma synchrony persisted in wild-type mice on Day 3. We did not observe significant changes for other frequency bands (beta: 15-25 Hz; high gamma: 50-70 Hz).

## DISCUSSION

Gamma-frequency sensory stimulation (GENUS) has been proposed as a therapy for restoring gamma synchrony, and has been shown to be effective for reducing neuropathology and producing beneficial behavioral effects in certain mouse models of Alzheimer’s disease. On the one hand, it is a simple, non-invasive and potentially inexpensive intervention that could have a transformative impact on a major human health challenge. On the other hand, the mechanism through which it operates remain mysterious. This makes it difficult to optimize, while also raising questions about how well it will translate beyond these particular mouse models and about whether it is acting to correct pathophysiological dysfunction vs. via indirect or nonspecific pathways. To address these issues, we studied the effects of GENUS in mice which have abnormalities in PV interneurons, gamma oscillations, and performance in a rule shifting task, and leveraged a method we had previously described for measuring cell type-specific deficits in gamma synchrony. GENUS rescued behavioral deficits in this model, but in contrast to 40 Hz optogenetic stimulation of PV interneurons and CLZ, the pro-cognitive effects of GENUS did not persist. Furthermore, unlike CLZ, GENUS did not restore normal patterns of synchrony.

From our previous studies, we know that we can detect increase in cross-hemispheric PV interneuron gamma synchrony in *Dlx5/6*^+/-^ mice. Furthermore, we know that in normal (wild-type) mice, cross-hemispheric PV interneuron gamma synchrony is necessary for rule shifts, i.e., disrupting it (optogenetically) prevents mice from learning rule shifts. Thus, the fact that we do not observe an increase in gamma synchrony after rule shift errors in *Dlx5/6*^+/-^ mice suggests that GENUS is acting through a mechanism other than restoring gamma synchrony between prefrontal PV interneurons. This is further supported by the observation that the pro-cognitive effects of GENUS, unlike those of synchronous optogenetic stimulation or CLZ, do not persist. One intriguing possibility is suggested by the observation that entrainment from sensory stimuli can spread across the cortex, demonstrated by mouse (Adaikkan et al., 2019) and human studies (Lerousseau et al., 2021). In the latter study, oscillatory activity recruited by sensory stimulation decayed rapidly (<1 second). Thus, gamma-frequency auditory stimulation may transiently entrain cortical activity, producing pro-cognitive effects, but may not do so via normal mechanisms that rely largely on PV interneurons. As a result, the entrainment may not be reflected in PV interneuron activity, particularly for mutant mice that have abnormal PV interneuron excitability, and as a result, may be able to elicit long-lasting changes in circuit function. An alternative is that GENUS elicits more nonspecific effects, e.g., changes in arousal, which transiently improve the performance of *Dlx5/6*^+/-^ mice during the period of stimulation.

In summary, these results underscore both the potential positive impact of GENUS on behavior and the importance of further studies to understand the exact mechanisms through which GENUS elicits these effects. It shows that even though rhythmic stimulation may entrain cortical activity, it may not consistently recruit endogenous cell types in a naturalistic manner, particularly in pathological conditions where the excitability of those cell types is altered. Future studies should explore whether augmenting this sensory stimulation, e.g., using multiple sensory modalities or drugs that target specific cell types of synapses, could overcome these limitations, and/or whether the effects we observed during rule shifting in *Dlx5/6*^+/-^ mice would translate to other models and tasks.

## ACKNOWLEDGEMENTS

Kathleen Cho provided advice and assistance with TEMPO analysis. This work was supported by a McKnight Memory and Cognitive Disorders award to VSS, and by NIH (R01MH121342 and R01NS116594 to VSS).

The authors declare no competing interests.

## MATERIALS AND METHODS

### Mice

All animal care and experiment procedures were in accordance with the guidelines of the National Institute of Health. Animal protocols were approved by the Administrative Panels on Laboratory Animal Care at the University of California, San Francisco. Prior to testing, mice were group-housed under normal lighting conditions in a 12-hour light/dark cycle. During testing mice were individually housed under a reversed light-dark cycle. Ad libitum food was provided until the beginning of the rule shifting task.

*Dlx5/6*^+/-^ mice were backcrossed to C57Bl/6 mice for at least six generations and crossed with PV-Cre line (Jackson Laboratory). Tail samples from litters were collected when they reached their weaning age. Genotyping was performed based on a previously described protocol (Wang et al., 2010). Both sexes were used for experiments and mice were 10-20 weeks old at the time of behavioral testing.

### Surgery

Mice were anesthetized using isoflurane (4% induction, 1.5% maintenance in 95% oxygen). Surgeries were performed on a stereotaxic instrument (Kopf Instruments). Before incision, mice were given subcutaneous buprenorphine and meloxicam, as well as lidocaine at the incision site. Doses were calculated based on mice’s weight. An incision above the skull was made to expose bregma and lambda for vertical alignment (difference < 0.05mm). A scalpel and 70% ethanol were used to score and clean the skull surface to improve dental cement adhesion. Craniotomy was performed to expose the brain surface above mPFC.

For dual-site TEMPO experiments, mice were injected bilaterally in the mPFC at three depths using the coordinates: 1.7 AP, ±0.3 ML, and -2.5, -2.25, and -2.0 DV. At each depth, we injected 0.2μl AAV1-CAG-DIO-Ace2N-4AA-mNeon (Virovek) and 0.1μl AAV2-Syn-tdTomato (Addgene). After viral injection, two fiber implants (Doric Lenses, MFC_400/430-0.48_2.8mm_ZF1.25_FLT) were slowly inserted into the brain at a 12-degree angle to 1.7 AP, ±0.76 ML, and -2.13 DV. Fiber implants were secured using C&B-Metabond (Parkell). The posterior portion of the incision site was sutured. Mice were allowed to recover for at least 5 weeks before beginning behavioral experiments.

### Rule shift task

We used the same task as previously described (Cho et al., 2020, 2015). Mice were singly housed and kept in a reverse light/dark cycle, and food-restricted to ∼85% of their ad libitum weight. For each trial, the mouse was placed in its home cage to explore two bowls, each with a digging medium and odor. Two types of digging media and two odors were used to form four possible cue combinations. The mouse could explore both bowls until choosing to dig in one. Digging was determined by the onset of continuous paw strokes which displaced media in one bowl. The correct bowl contained a buried peanut butter chip food reward. If the mouse dug in the correct bowl, it was allowed to dig until the reward was found. Once the mouse began to dig in the incorrect bowl, the other (correct) bowl was removed. The mouse was allowed to explore the incorrect bowl until it stopped digging. Following each trial, the mouse was transferred to a holding cage. After correct trials, the mouse stayed in the holding cage for 10-20 seconds until the onset of the next trial. Following incorrect trials, the mouse received a time-out of 2 minutes in the holding cage, during which a translucent lid was placed over the top of the holding cage. For both the initial association and rule shift portions of the task, the mouse was required to reach a criterion of 8 correct out of 10 consecutive trials. We further required that of these 8 correct trials, the fraction of choices corresponding to a particular cue combination should >62.5% (5/8 trials). Additional trials were added if these criteria were not met.

For GENUS auditory stimulation, the 40 Hz train of sound clicks started playing after the mouse reached the criterion for the initial association portion of the task. We added 3 additional initial association trials after the mouse met the learning criterion, to allow the mouse to habituate to the clicks. Clicks were played throughout the rule shift portion of the task until the mouse reached the learning criterion at which point testing ended.

We categorized errors during the rule shift part of the task as *perseverative* or *random*. Perseverative errors are trials in which the mouse dug in the bowl that would have been correct based on the initial association rule. On random error trials, the mouse digs in a bowl that would be incorrect based on both the current and previous rule.

### GENUS auditory stimulation

We used the previously described protocol (Martorell et al., 2019). A speaker set was placed out of reach and directly above the mouse home cage. We utilized a MATLAB script pure tone generator created by Kamil Wojcicki (https://www.mathworks.com/matlabcentral/fileexchange/34058-pure-tone-generator). We wrote a script to play a 10 kHz pure tone at 40 Hz with a sampling frequency of 88.2 kHz to generate the 40 Hz sound clicks. The script is written to play a 20-second train of sound clicks, and repeated 7,200 times, with a theoretical duration of 144,000 seconds. Recording usually finishes before the end of the script duration. The tones are delivered at 60 dB, measured by Decibel X app on the same level as the mouse home cage. Sound clicks were played continuously during the rule shift portion of the task.

### Voltage photometry (TEMPO)

Bulk fluorescence signals from each recording site were measured using TEMPO (Marshall et al., 2016), following our previously-published protocol (Cho et al., 2020).

### Optical apparatus

Fiber-optic cannulas implanted in the brain region were connected with a fiber-optic patch cord (Doric Lenses, MFP_400/430/900-0.57_2m_FC-ZF1.25(F)_LAF) to a mini-cube (Doric Lenses, FMC5_E1(460-490)_F1(500-540)_E2(555-570)_F2(580-680)_S). The fiber was connected to the 1.25mm implant via a zirconia sleeve with the corresponding diameter and cleaned using 70% ethanol before each recording session.

The mini-cube creates a light path to monitor two different fluorophores. The dichroic mirror and cleanup filters were chosen to match the excitation and emission spectra of Ace-mNeon and tdTomato (Ace-mNeon channel: excitation, 460-490 nm, emission, 500-540 nm; tdTomato Channel: excitation, 555-570 nm, emission, 580-680 nm). All five ports on the mini-cube were connected with respective FC optic connections (one to the animal, two excitation lines from LEDs, and two emission lines to photodetectors).

Excitation for both channels was provided by LEDs (Thorlabs, M490F3 and M565F3), controlled by an LED controller (Thorlabs, DC4104). LEDs were connected to the mini-cube through a fiber-coupled patch cord (Thorlabs, M75L01). The emission channels from the mini-cube were connected to individual photoreceivers (Femto, OE-200-Si-FC; bandwidth 7 kHz, AC coupled, ‘Low’ gain) using a large-core high-NA fiber (Doric Lenses, MFP_600/6300/LWMJ-0.48_0.5m_FC-FC).

### Lock-in modulation and detection

Each of the four channels of excitation LEDs was sinusoidally modulated at a distinct frequency to minimize potential crosstalk between fluorophores and hemispheres. The emission channels were then demodulated using the same frequency. Both modulation and demodulation was performed by a lock-in amplifier (Stanford Research Systems, SR860). The following frequencies were selected for each channel: Ace-mNeon side 1: 3 kHz; Ace-mNeon side 2: 2.5 kHz; tdTomato side 1: 4 kHz; tdTomato side 2: 3.5 kHz. Low-pass filters were selected to reject noise (attenuation of signals were approximately -1 dB at 20 Hz, -3 dB at 40 Hz and -6 dB at 60 Hz).

### TEMPO recording

The analog signal from the lock-in amplifier was digitized using a processor (Tucker-Davis Technologies, RX-8). Recording software Synapse (Tucker-Davis Technologies) on a PC was used to record the data stream and time-locked video from a generic infrared USB webcam (Ailipu Technology, ELP-USC100W05MT-DL36). The data stream was sampled at 3 kHz.

### Histology and imaging

Mice were injected with 0.1ml Euthasol and transcardially perfused with 0.01M phosphate base saline (PBS) and 4% paraformaldehyde (PFA) in PBS. Brains were post-fixed in 4% PFA for 24-48 hours before being rinsed and stored in PBS. Brains were sliced at 70-μm thickness using a Leica VT1000S Vibratome. Imaging was performed using Nixon Eclipse 90i. We verified virus expression and fiber placement for all mice.

### TEMPO analysis

We used custom MATLAB (Mathworks) scripts and the signal processing toolbox to analyze the TEMPO recordings. Our method to extract signal used linear regression and bootstrapping to measure the number timepoints that were superior to shuffled data for predicting zero-phase-lag cross-hemispheric synchronization, as previously described (Cho et al., 2020). All four signals were filtered around the frequency of interest. A linear regression analysis was used to predict the right mNeon signal using left mNeon, left tdTomato, and right tdTomato signals. The goodness of fit was compared to regression using shuffled left mNeon signal. *R*^*2*^ values were calculated using 1-s segments and compared to the 99^th^ percentile of the *R*^*2*^ values from shuffled data to measure the zero-phase-lag synchronization between the left and right mNeon signals. We compared the *R*^*2*^ values smoothed over a 5-min time window after the start of digging for the first 5 rule shift trials. Correct and error trial outcomes were compared. All mice made both correct and error outcomes during the first five rule shift trials.

### Data analysis and statistics

Statistical analyses were performed using Prism 7 (GraphPad). Quantitative data were expressed as mean, and error bars represent the s.e.m. Group comparisons were made using one-way repeated-measure ANOVA followed by Tukey’s post hoc tests for multiple comparison. Paired and unpaired two-tail Student’s t-tests were used to make the single-variable comparisons. P values were symbolled as follow: *, <0.05; **, <0.01; ***, <0.001; ****, <0.0001. Comparison with no asterisk had P >0.05 for no significance.

